# Role of the *osaA* transcription factor gene in development, secondary metabolism and virulence in the mycotoxigenic fungus *Aspergil lus flavus*

**DOI:** 10.1101/2025.11.26.690797

**Authors:** Farzana Ehetasum Hossain, Apoorva Dabholkar, Jessica M. Lohmar, Matthew D. Lebar, Brian M. Mack, Ana M. Calvo

## Abstract

*Aspergillus flavus* colonizes oil-seed crops contaminating them with aflatoxins, highly carcinogenic mycotoxins that cause severe health and economic losses. Genetic studies may reveal new targets for effective control strategies. Here we characterized a putative WOPR transcription factor gene, *osaA*, in *A. flavus*. Our results revealed that *osaA* regulates conidiation and sclerotial formation. Importantly, deletion of *osaA* reduces aflatoxin B_1_ production, while, unexpectedly, transcriptome analysis indicated upregulation of aflatoxin biosynthetic genes, suggesting post-transcriptional or cofactor-mediated regulation. Cyclopiazonic acid production also decreased in absence of *osaA*. In addition, the *osaA* mutant exhibited upregulation of genes in the imizoquin and aspirochlorine clusters. Moreover, *osaA* is indispensable for normal seed colonization; deletion of *osaA* significantly reduced fungal burden in corn kernels. Aflatoxin content in seeds also decreased in the absence of *osaA*. Furthermore, deletion of *osaA* caused a reduction in cell-wall chitin content, as well as alterations in oxidative stress sensitivity, which could in part contribute to the observed reduction in pathogenicity. Additionally, promoter analysis of *osaA*-dependent genes indicated potential interactions with stress-responsive regulators, indicated by an enrichment in Sko1 and Cst6 binding motifs. Understanding the *osaA* regulatory scope provides insight into fungal biology and identifies potential targets for controlling aflatoxin contamination and pathogenicity.

**Key Contribution:** *Aspergillus flavus osaA* controls morphological and chemical development, as well as phytopathogenicity, and could be a promising target for a control strategy against *A. flavus* to reduce health risks and economic losses associated with aflatoxin contamination.

## 1. Introduction

*Aspergillus flavus* is a ubiquitous filamentous fungus present in soil and decaying organic matter. It disseminates primarily through the formation of asexual spores called conidia, which are carried by wind, insects, and rain [1,2]. These conidia are highly adaptable and capable of germinating under a wide range of environmental conditions, and the resulting mycelial growth has been reported across temperatures spanning from 12°C to 50°C [3]. In addition to conidia, *A. flavus* produces sclerotia to persist in adverse environmental conditions [1,4]. Both conidia and sclerotia are important structures contributing to fungal dissemination and resilience [5].

From an ecological and agricultural perspective, *A. flavus* is particularly notorious as an opportunistic plant pathogen that colonizes oilseed crops such as corn, peanuts, cotton, sorghum, and tree nuts, during both pre- and post-harvest [6,7]. In peanuts and cotton, dispersal occurs mainly through rain and soil, while in corn, infection typically takes place via silk colonization and kernel invasion by fungal mycelia [1]. The pathogen’s broad host range, environmental adaptability, and persistence mechanisms make it a formidable challenge for agricultural systems worldwide.

In addition to its agricultural impact, *A. flavus* poses a serious clinical threat, especially in immunocompromised individuals. It is the second leading cause of invasive aspergillosis (IA) after *Aspergillus fumigatus*, contributing to pulmonary infections and systemic mycoses in patients undergoing chemotherapy, organ transplantation, or corticosteroid therapy [2]. This dual threat to plants and human health underscores the urgency of developing effective control strategies.

A major concern associated with *A. flavus* plant colonization is its ability to synthesize aflatoxins. Of these, aflatoxin B_1_ (AFB_1_) is the most toxic and has been classified by the World Health Organization as a Group 1A human carcinogen [8]. In addition to aflatoxins, *A. flavus* produces other secondary metabolites such as cyclopiazonic acid (CPA) and aflatrem [9,10]. Aflatoxin B_1_ (AFB_1_) is found contaminating crops for human consumption as well as in animal feed, and consequently in animal-derived products, posing a serious threat to both animal and human health. [8]. In humans, aflatoxin-contaminated food can cause acute symptoms such as jaundice, abdominal pain, and liver failure, with outbreaks reaching fatality rates up to 40%. [11,12]. Chronic exposure leads to immune suppression, growth stunting, and higher risks of cancers, particularly hepatocellular carcinoma, and increases susceptibility to infectious diseases like HIV/AIDS. [11,13–15].

Beyond its severe health consequences, aflatoxin contamination also leads to significant economic losses, particularly in the United States and other developed countries. In the United States alone, aflatoxin contamination of corn results in annual losses ranging from USD $52 million to $1.68 billion, especially during years of drought and high heat [16]. The U.S. peanut industry lost $52.28 per metric ton in 2019, the worst aflatoxin year in recent history [17]. According to Farm Progress report, aflatoxin costs the U.S. peanut industry as much as $126 million each year [18]. Developed countries impose stringent regulatory limits on aflatoxin levels in food and feed products. While essential for safeguarding public health, these regulations can result in substantial economic losses when contamination exceeds the allowable thresholds [4]. Globally, it is estimated that approximately 25% of the world’s food crops are lost each year due to aflatoxin contamination [19]. Additional economic burdens arise from rejected agricultural exports, livestock losses, and the costs of testing and monitoring for aflatoxins [6]. Aflatoxin contamination costs Africa over USD 750 million annually, while compliance with EU aflatoxin regulations is estimated to cost food exporters USD 670 million each year, highlighting the substantial economic burden [20]. In contrast, many developing nations lack regulatory frameworks or effective enforcement mechanisms, resulting in higher rates of aflatoxin exposure. As a consequence, an estimated 4.5 billion people worldwide are at risk of chronic aflatoxin ingestion [21].

Efforts to manage *A. flavus* colonization and toxin contamination have met limited success. Current strategies, including chemical fungicides (e.g., azoles), biological control, and breeding for host resistance, often fall short due to variability in efficacy, environmental resistance, and evolving fungal populations [22]. Notably, the widespread use of azole fungicides in agriculture is believed to contribute to the emergence of azole-resistant fungal strains, thereby compromising both crop protection and clinical antifungal therapy [23]. These limitations necessitate the discovery of novel genetic targets and regulatory mechanisms to control the growth, development, mycotoxin production and pathogenicity of *A. flavus*. A promising approach is the exploration of transcriptional regulators that could control developmental and metabolic pathways and possibly virulence. Among these, Gti1/Pac2 domain (also known as the WOPR domain) containing proteins are highly conserved fungal-specific transcriptional regulators that play a central role in controlling cell morphology, development, secondary metabolite production, and pathogenesis across diverse fungal species [24–26]. The WOPR domain has been shown to be important for DNA-protein interactions and contains two globular domains that are referred to as WOPRa and WOPRb [26,27]. This class of transcriptional regulators are typically separated into two main groups in fungi which are named Gti1/Wor1/Ryp1/Mit1/Rge1/Ros1/Sge1 or Pac2 with several members of these groups having been functionally characterized in a variety of fungal species [26]. The name “WOPR” is derived from a group of functionally related transcription factors: Wor1 in *Candida albicans*, Pac2 in *Schizosaccharomyces pombe*, and Ryp1 in *Histoplasma capsulatum*, each of which is critical for development and virulence [25]. *Candida albicans*, Wor1 orchestrates the white-to-opaque phenotypic switch, a key process that regulates mating competence and influences tissue specificity during infection [28]. In *Schizosacchoarymces pombe*, Pac2 is thought to mediate nutritional signaling that activates *ste11*, a transcription factor essential for sexual development where as Gti1 was shown to function downstream of the Wis1-Sty1 MAPK pathway and to be essential for gluconate absorption [29,30]. In *Histoplasma capsulatum*, Ryp1 governs the dimorphic switch between yeast and mycelial forms and regulates the expression of hundreds of genes required for pathogenicity [31]. Notably, the best-characterized WOPR protein, Wor1 from *C. albicans*, binds to a conserved 9-nucleotide DNA motif (TTAAAGTTT). This motif is also recognized by its orthologs Ryp1 in *H. capsulatum* and Mit1 in *Saccharomyces cerevisiae*, reflecting the remarkable evolutionary conservation of WOPR domain–DNA interactions over an estimated 600 million to 1.2 billion years of fungal divergence [25,32].

In the model fungus *Aspergillus nidulans*, OsaA, a predicted WOPR domain-containing protein, was identified as a downstream regulator of the global regulator VeA, a member of the velvet complex that governs fungal development and secondary metabolism. OsaA contributes to the balance between asexual and sexual morphogenesis [33]. A more recent study in *A. fumigatus* demonstrated that the OsaA homolog plays a crucial role in regulating colony growth, conidiation, secondary metabolite production, and virulence in mouse models [34]. OsaA-deficient strains in *A. fumigatus* also showed impaired thermotolerance, reduced cell wall stability, and hypersensitivity to oxidative stress, traits essential for survival and infection [34].

Interestingly, *A. flavus* possesses an OsaA homolog that shares 68% sequence identity with its *A. nidulans* counterpart [33], but its role in this agriculturally relevant species remains uncharacterized. Given the established role of Gti1/Pac2 (WOPR) proteins in other fungi, we hypothesized that *osaA* functions as a major regulator in *A. flavus*, integrating development, mycotoxin biosynthesis and virulence.

In the present study, we used a combination of genetic and transcriptomic analyses, as well as biochemical approaches and seed colonization bioassays to characterize the function of *osaA* in *A. flavus*. Our findings establish *osaA* as an important regulatory node in this fungus and highlight its potential as a molecular target for controlling fungal growth, aflatoxin production and virulence, offering new avenues for integrated disease and toxin management in agriculture and health.

## 2. Results

### 2.1. OsaA in A. flavus contains a Gti/Pac2 (WOPR) domain which is conserved with orthologs in other Aspergillus species and other species in the Ascomycota phylum

The amino acid sequence of the *A. flavus* OsaA was obtained from FungiDB (accession number: F9C07_2071416). OsaA orthologous proteins were found by using NCBI BLASTP. Our phylogenetic analysis revealed that OsaA in *Aspergillus spp*. is conserved with other Gti1/Pac2 (WOPR) proteins in fungi but is most closely related to Gti1 proteins (Figure 1). Additionally, *A. flavus* OsaA and orthologous proteins contain a conserved WOPR domain in their N-terminal region. WOPR-domain proteins are a fungal-specific family of transcriptional factors that are involved in multiple biological processes [24]. Specifically, the predicted OsaA amino acid sequence contains the two-conserved WOPR subdomains (12-87 residues and 166-187 residues), separated by a characteristic less conserved linker (Figure S1).

**Figure 1.**
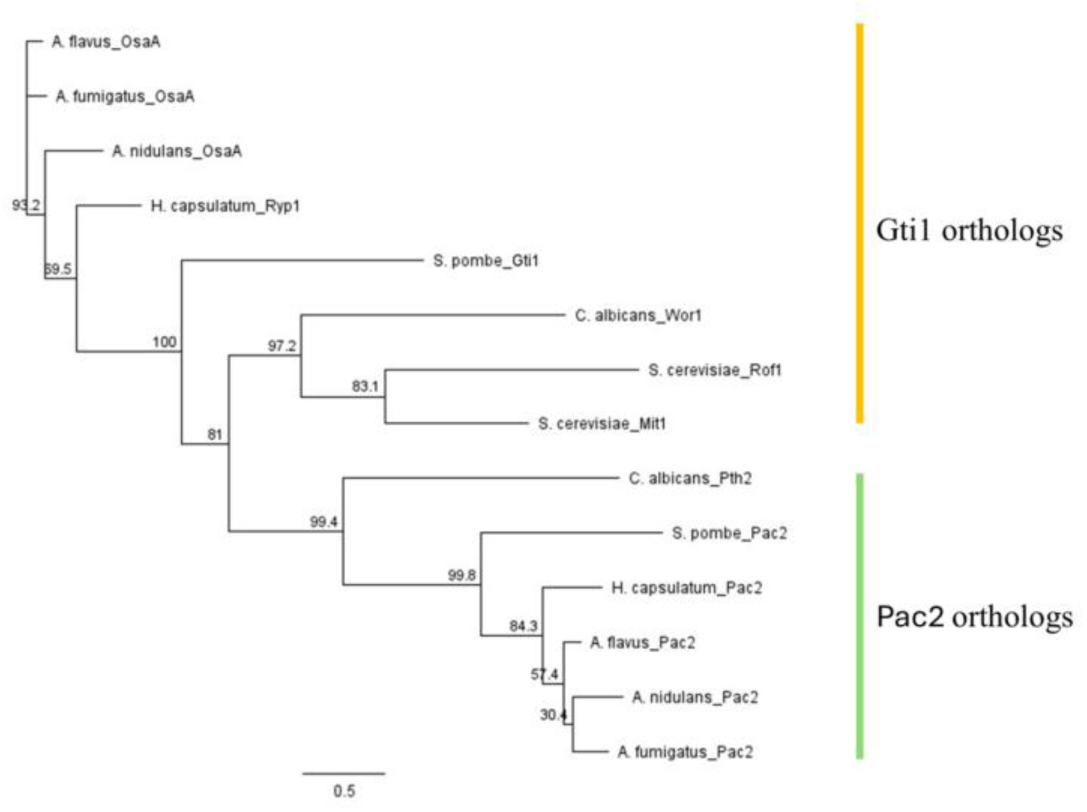
Maximum likelihood phylogenetic tree of Gti1 and Pac2 proteins from *Aspergillus* species and phylum Ascomycota. Construction of a maximum likelihood phylogenetic tree was carried out in Geneious software with the PhyML plugin (version 3.3.20180621) with a LG substitution model and 1,000 bootstrap replicates.

### 2.2. osaA regulates development in A. flavus

Recently it was shown that *osaA* plays a significant role in morphogenesis of the opportunistic human pathogen *A. fumigatus* [34]. Moreover, in the model fungus *A. nidulans osaA* regulates development primarily by suppressing formation of sexual structures [33]. To understand the role of *osaA* in the agriculturally relevant fungus *A. flavus*, *osaA* deletion strain (TMR1.1) and *osaA* complementation strain (TFEH 8.1) were constructed by previously described methods [35,36]. The *osaA* deletion strain (Δ*osaA*), where the *osaA* gene was replaced with the *pyrG* marker, was confirmed by PCR, yielding the expected 3.37 kb PCR product (Figure 2A, C). The complementation strain (Com), where the *osaA* allele was reintroduced into the *osaA* mutant, was also confirmed by PCR, obtaining a 3.304 kb target size DNA band (Figure 2B, D). Expression analysis of *osaA* in the wild type, deletion, and complementation strains was carried out by qRT-PCR. As expected, the Δ*osaA* lost *osaA* expression, while the complementation strain presented *osaA* expression levels similar to that of the wild type (Figure 2E).

**Figure 2.**
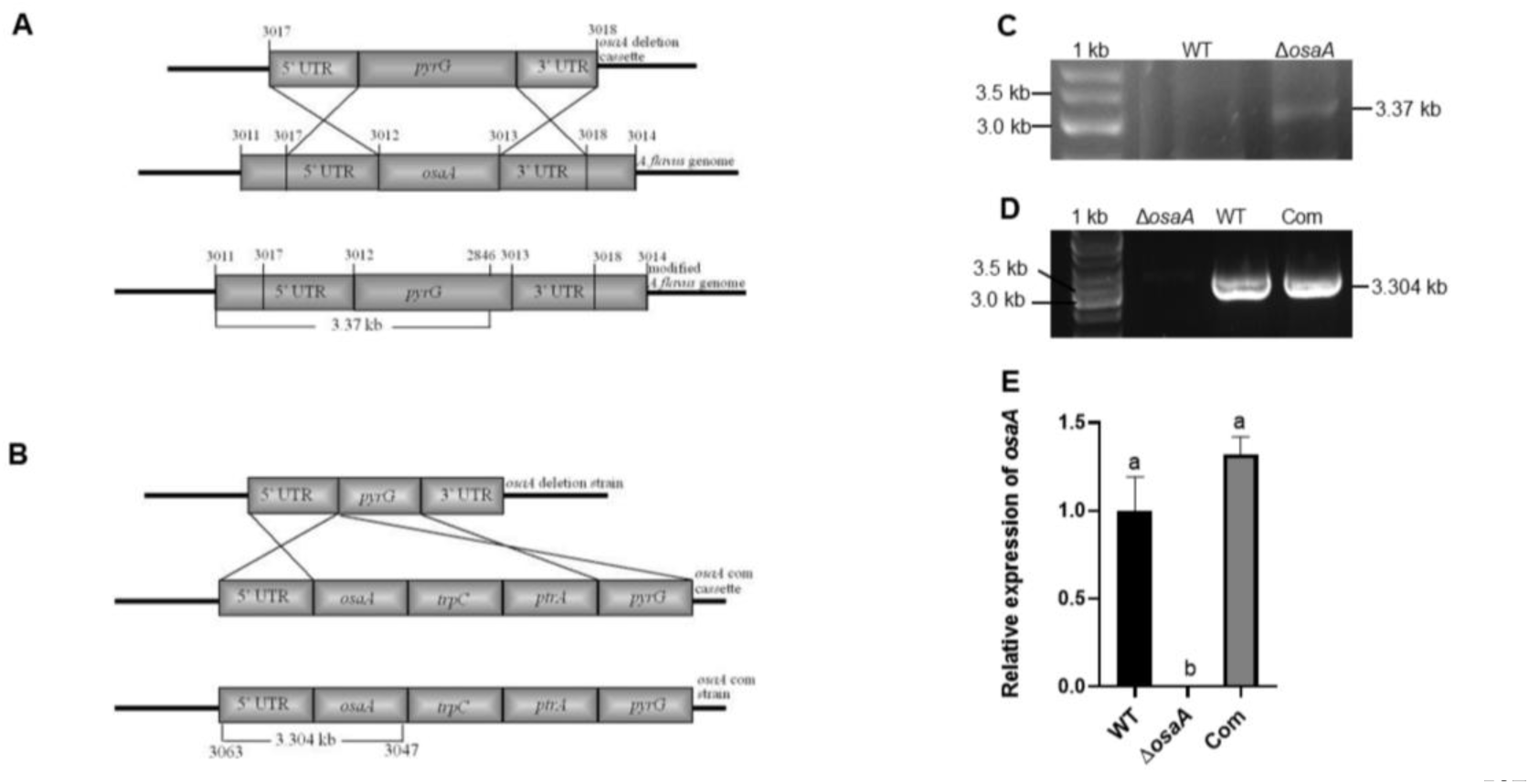
Generation and confirmation of Δ*osaA* and complementation strains. **(A)** Diagram showing the replacement of *osaA* with the *pyrG* marker by a double-crossover event. **(C)** Confirmation of the Δ*osaA* strain by diagnostic PCR using primer 3011 and 963 (Table S1). The expected band size was 3.37 kb, wild-type strain was used as control. **(B)** Representation of the complementation cassette and its insertion into the Δ*osaA* strain at the same locus **(D)** Results of PCR, confirming the integration of the complementation fusion cassette carrying the *osaA* wild-type allele in the Δ*osaA* strain, using primers 3065 and 3047(Table S1). The expected band size was 3.304 kb, wild type and *osaA* mutant strains were used as positive and negative controls, respectively. **(E)** Expression analysis of *osaA* by qRT-PCR using primer 3071 and 3072 (Table S1). The relative expression was calculated using the 2^−ΔΔ*CT*^ method, as described by [74]. The expression of 18S rRNA was used as an internal reference. Values were normalized to the expression levels in the wild type, considered as one. Error bars represent the standard errors. Different letters on the columns indicate values that are statistically different (P < 0.05).

In present study, we observed that colony diameter was reduced (22.05% and 14.11% respectively) in Δ*osaA* compared to the wild-type control on both day 5 and day 7 (Figure 3A, B, C), indicating that *osaA* is required for normal growth in *A. flavus*. Our study also showed that conidial production was significantly increased in the *osaA* deletion strain compared to the control at all time-points analyzed (Figure 3D, E), revealing the role of *osaA* as a repressor of conidiation in this fungus.

**Figure 3.**
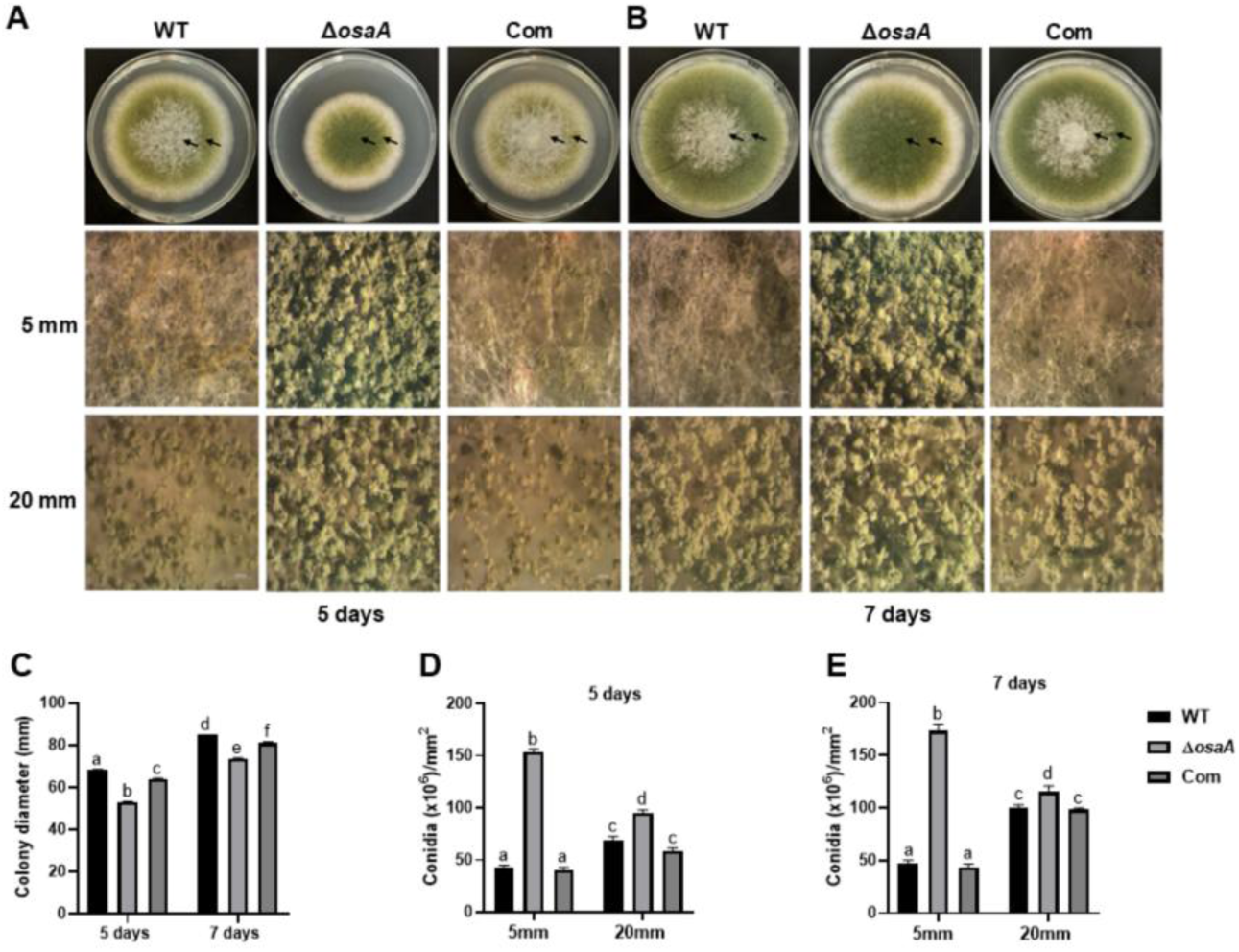
osaA affects colony growth and conidiation in A. flavus. Wild type (WT), deletion *osaA* (Δ*osaA)* and complementation *osaA* (Com) were point-inoculated on PDA medium at 30°C. **(A)** Photographs of colonies taken after five days of incubation. Here, two arrows correspond to 5 mm and 20 mm away from the colony center. The experiment was done in triplicates. Second and third rows show closeup plate image of conidiophores under a Leica dissecting scope at 32X magnification, 5 mm and 20 mm away from the colony center, respectively. **(B)** Photographs taken after seven days of incubation. Here, two arrows correspond to 5 mm and 20 mm away from the colony center. Second and third rows are closeup plate image of conidiophores at 32X magnification of 5 mm and 20 mm away from the colony center, respectively. **(C)** Colony diameter on PDA medium after 5 and 7 days at 30°C. **(D)** Conidial quantification on PDA medium after 5 days at 30°C. Counting was carried out 5 mm and 20 mm away from the colony center. **(E)** Conidial quantification on PDA medium after 7 days at 30°C. Conidial count was carried out 5 mm and 20 mm away from the colony center. The error bars represent standard error. Different letters on the columns indicate values that are statistically different (P < 0.05), as determined by two-way ANOVA with Tukey test comparison.

In addition, *osaA* also influences sclerotial formation in *A. flavus*. Specifically, we found that *osaA* is essential for normal sclerotial development. In point-inoculated cultures, *osaA* deletion strains produced significantly less sclerotia than the controls on day 14, while in top-agar cultures, where crowdedness is a factor, sclerotial formation was completely abolished in the mutant (Figure 4).

**Figure 4.**
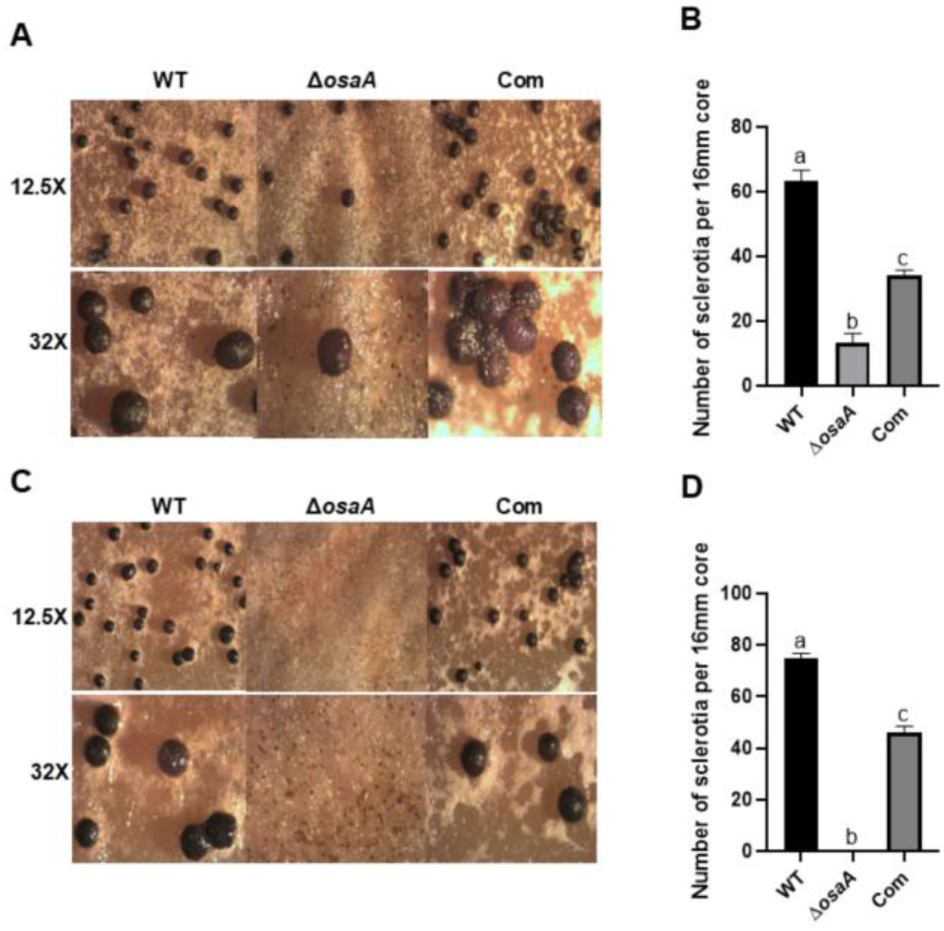
osaA positively regulates sclerotial formation. **(A)** Wild type (WT), deletion *osaA* (Δ*osaA)* and complementation *osaA* (Com) were point-inoculated on Wickerham medium and grown at 30°C in the dark for 14 days. Sclerotia were photographed after spraying the plate with 70% ethanol to remove conidia, visualizing them under a dissecting microscope at 12.5X and 32X magnification. **(B)** Sclerotial quantification of Wickerham cultures after 14 days. One 16 mm diameter core was collected 5 mm away from the center and sclerotia were counted. **(C)** Wild type, deletion *osaA* and complementation *osaA* were top agar inoculated (10^6^ spores/ml) on Wickerham medium and grown at 30°C in dark conditions for 14 days. Sclerotia were photographed after spraying the plate with 70% ethanol to remove conidia, observing them under a dissecting microscope at 12.5X and 32X magnification. **(D)** Sclerotial quantification on Wickerham medium after 14 days. One 16 mm diameter core was collected 5 mm away from the center and sclerotia were counted. Error bars represent the standard error. Columns with different letters represent values that are statistically different (p< 0.05).

### 2.3. Secondary metabolism is influenced by osaA

Development and secondary metabolite production are genetically linked processes in *Aspergillus* species [5]. In *A. fumigatus,* secondary metabolite production was influenced by *osaA* [34]. Therefore, it is possible that *osaA* also regulates the production of secondary metabolites in *A. flavus*. In this study we evaluated the role of *osaA* in regulating biosynthesis of aflatoxin B_1_ and CPA in *A. flavus.* Our TLC analysis showed that AFB_1_ production dramatically decreased in the *osaA* deletion strain compared to the wild type (Figure 5A). Densitometry of TLC band intensity indicated that such reduction was approximately 95.1% with respect to the wild-type control (Figure 5B). Additional LC-MS analysis also confirmed that AFB_1_ levels were significantly reduced, approximately 84.62% in the absence of *osaA* with respect to the wild type (Figure 5C), supporting that *osaA* positively regulates the biosynthesis of AFB_1_. Moreover, the LC-MS analysis also showed that production of CPA significantly decreased approximately 54.65% in the *osaA* deletion mutant compared to the control (Figure 5D), suggesting that *osaA* positively regulates biosynthesis of CPA. Complementation of the deletion mutant with the *osaA* wild-type allele rescued AFB_1_ and CPA production to levels similar to those in the wild type.

**Figure 5.**
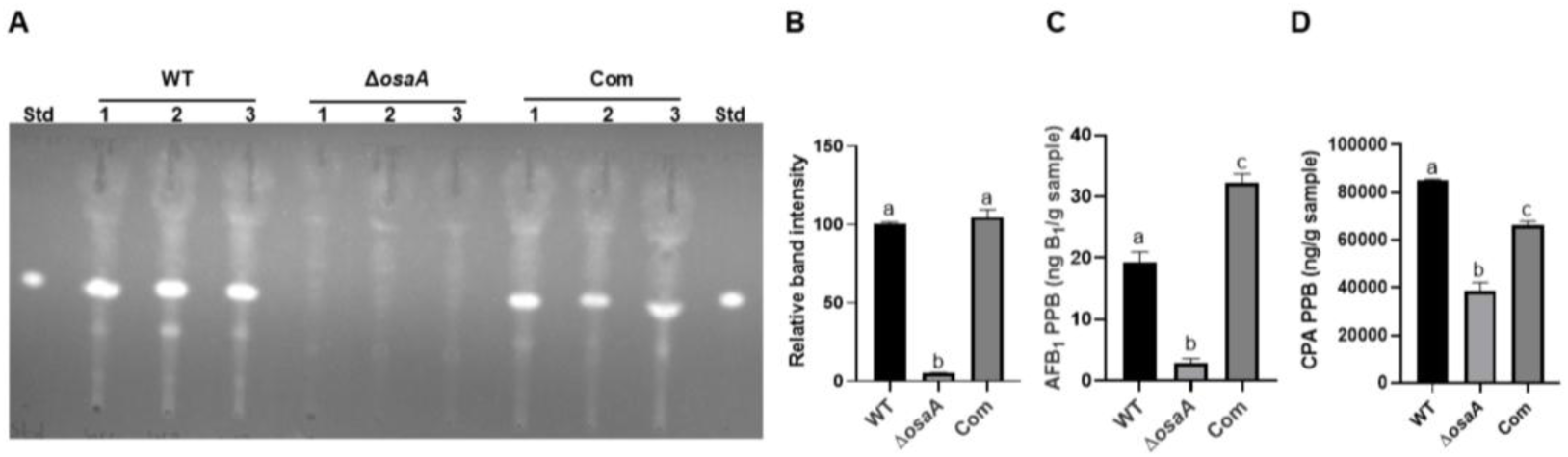
osaA positively influences A. flavus AFB_1_ and CPA production. **(A)** TLC results of wild type (WT), deletion *osaA* (Δ*osaA)* and complementation *osaA* (Com) point-inoculated on PDA medium and incubated at 30°C under dark conditions for 7 days. Three 16 mm diameter cores were collected 5 mm from the colony center and toxin was extracted using chloroform. Toluene: ethyl acetate: formic acid (50:40:10) solvent system was used. Aflatoxin B_1_ was used as Standard. **(B)** Densitometry for corresponding AFB_1_ band intensity of TLC image in (A), carried out with GelQuant. **(C)** LC-MS results of AFB1 content in extracts used in (A). **(D)** LC-MS results of CPA content in extracts used in (A). Error bars represent standard error. Different letters on columns represent values that are statistically different, P<0.05.

### 2.4. osaA affects sensitivity to environmental stresses in A. flavus

Fungi need to adapt to survive environmental stresses. It has been shown that *osaA* influences *A. fumigatus* to resist high temperature and oxidative stress [34]. Our results show that *A. flavus osaA* also plays a role in temperature and oxidative stress sensitivity. Fungal strains were grown at a range of temperatures, 25°C, 30°C, 37°C and 42°C (Figure S2). Percentages of growth rate change were calculated with respect to the colony diameter observed at 30°C. In the absence of *osaA*, colony growth reduction was significantly higher than in the controls at 25°C, indicating that *osaA* is relevant for cold resistance. In addition, all the strains tested showed the highest growth reduction at 42°C. Interestingly, Δ*osaA* slightly recover colony growth at 37°C.

With respect to sensitivity to oxidative stress, Δ*osaA* showed an increase in tolerance to menadione with respect to the control strains at 0.6 mM, a menadione concentration where the control strains were unable to grow (Figure S3A). On the other hand, the Δ*osaA* strain presented increased sensitivity to H_2_O_2_ with respect to the controls (Figure S3B); Δ*osaA* colonies showed a further colony growth reduction at 0.3% with respect to the wild type and complementation strains, and growth was completely abolished at 0.4% in the absence of *osaA*.

### 2.5. osaA is necessary for normal cell-wall chitin content in A. flavus

Based on the observed *osaA*-dependent change to environmental stress sensitivity we hypothesized that they could be, at least in part, due to alterations in cell wall composition in the Δ*osaA* strain. Fungal cell walls are a crucial structure providing support, protection, and regulating interactions with the environment. In *A. fumigatus* cell wall stability was also decreased in the absence of the *osaA* homolog [34]. Our cell wall composition analysis revealed that chitin content in *A. flavus osaA* deletion strain was significantly decreased, 38.39%, compared to the control strains (Figure S4).

### 2.6. osaA is indispensable for A. flavus seed infection

Previously, it was shown that *A. fumigatus osaA* was important in the infection of immunocompromised animals [34]. Based on this and the fact that *osaA* affects multiple aspects of *A. flavus* biology, it is likely that *osaA* could also have a role in plant seed infection and colonization. To test this hypothesis, viable B73 corn seeds were infected with the wild type, Δ*osaA* and *osaA* complementation strain. Cultures were photographed after 7 days of incubation (Figure 6A). In this experiment, levels of ergosterol were used as an indicator of fungal burden present in the infected plant tissue (Figure 6B). Seeds infected with the Δ*osaA* mutant strain contained significantly less ergosterol than seeds infected with the control strains. Importantly, the absence of *osaA* resulted in a statistically significant decrease in AFB_1_ and AFB_2_ production in viable seeds infected with the Δ*osaA* strain compared with the controls (Figure 6C, 6D).

**Figure 6.**
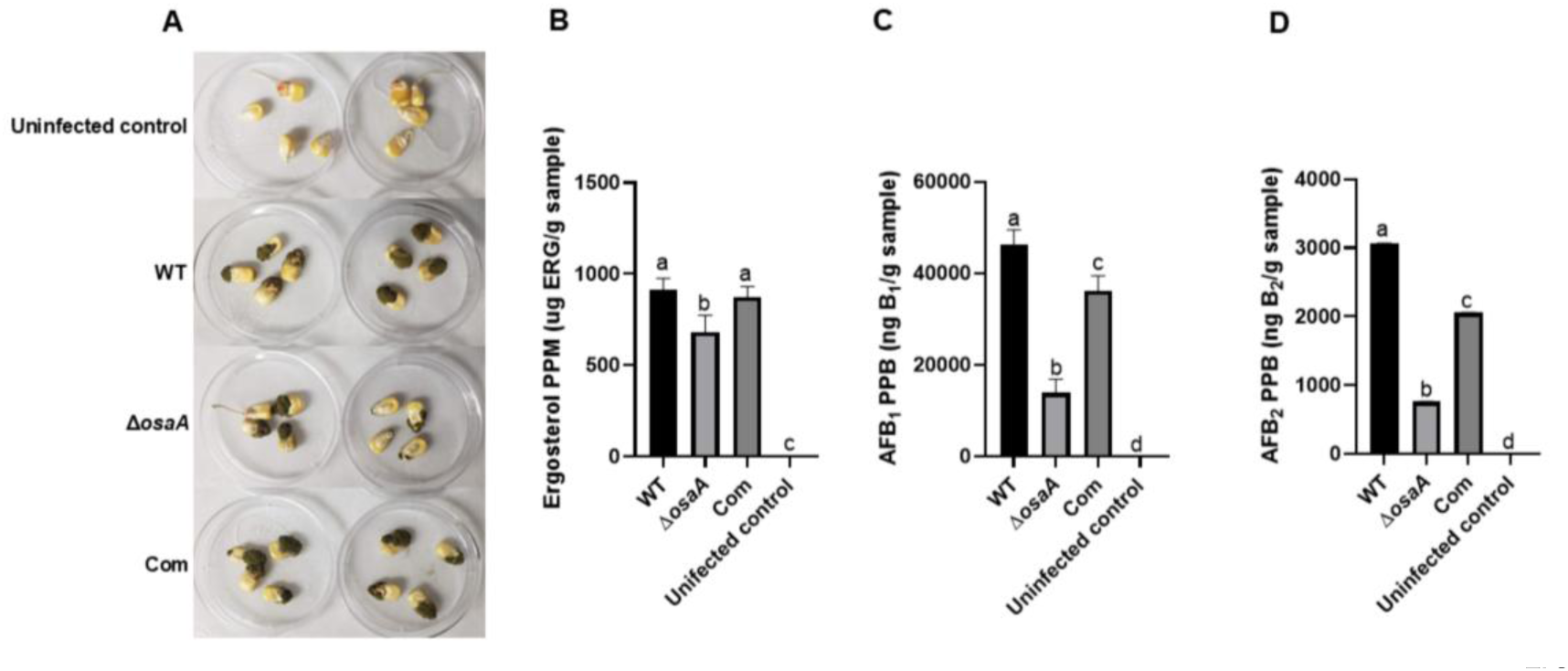
*osaA* is required for normal pathogenicity during *A. flavus* infection of corn seeds. **(A)** Wild type (WT), deletion *osaA* (Δ*osaA)* and complementation *osaA* (Com) were inoculated on the corn seeds and incubated 7 days. LC-MS quantification of fungal ergosterol **(B)**, aflatoxin B_1_ **(C)** and aflatoxin B_2_ **(D)** content in the infected seeds after 7 days of incubation. Error bars represent the standard error. Columns with different letters represent values that are statistically different (P < 0.05).

### 2.7. Global transcriptional changes induced by osaA deletion

RNA-Seq analysis was performed to investigate the *osaA*-dependent transcriptome in *A. flavus*. Our analysis revealed substantial transcriptome alterations in the Δ*osaA* mutant, with 1,488 differentially expressed genes (DEGs) compared to the wild type; 546 upregulated and 942 downregulated genes (Figure 7). Volcano plots indicated that most transcriptional changes were directly attributable to the absence of *osaA* in the mutant.

**Figure 7.**
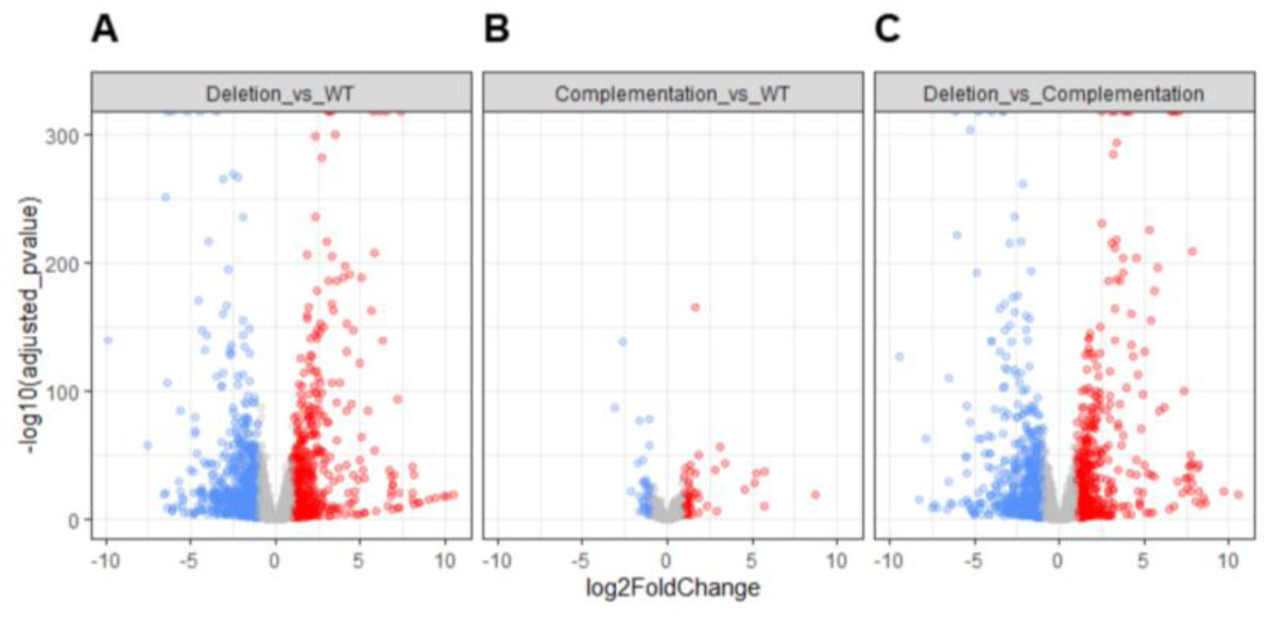
Volcano plots displaying differentially expressed genes (DEGs). **(A)** Deletion vs. Wild type; **(B)** Complementation vs. Wild type and **(C)** Deletion vs. Complementation. The *y*-axis represents the negative log10 of the adjusted *p* value of the differential expression results and the x-axis represents the log2 fold change value. The significantly up- and down-regulated genes are shown as red dots and blue dots, respectively (*p* < 0.05, fold change >2). Grey indicates non-significant genes that did not cross the threshold of adjusted p value or fold-change.

### 2.8. Gene Ontology and Heatmap analyses results

#### 2.8.1. Downregulated genes

Gene Ontology (GO) and functional enrichment analyses of downregulated genes in the Δ*osaA* mutant revealed significant enrichment of several biological categories (Table S3). Most notably, genes associated with the apoplast (96 out of 470 genes, adjusted p-value < 4e-14, odds ratio = 3.0) and apoplastic effectors (38 out of 146 genes, adjusted p-value < 8e-8, odds ratio = 3.9) were significantly downregulated. Additionally, genes related to membrane components (GO:0016020; 224 out of 1,953 genes, adjusted p-value = 2.8e-5, odds ratio = 1.5) and extracellular regions (GO:0005576; 27 out of 127 genes, adjusted p-value = 9.5e-4, odds ratio = 3.0) showed reduced expression. Cytoplasmic effectors were also downregulated (19 out of 92 genes, adjusted p-value = 0.021, odds ratio = 2.9). Other significantly downregulated functional categories included cutinase activity (GO:0050525; 4 out of 5 genes, adjusted p-value = 0.025, odds ratio = 44.2) and the pentose phosphate pathway (9 out of 28 genes, adjusted p-value = 0.029, odds ratio = 5.2) (Table S3).

#### 2.8.2. Upregulated genes

Strikingly, our analysis revealed a strong enrichment of secondary metabolism genes among up-regulated DEGs in the Δ*osaA* (Table S3), except two of the clustered aflatrem genes, that were downregulated (Table S2). Most prominently, genes belonging to the aflatoxin biosynthetic cluster (SMURF cluster 54; 26 out of 30 genes, adjusted p-value < 1.9e-7, odds ratio = 134.2) were highly upregulated (Table S3, Figure 8). Intriguingly, this transcriptional upregulation of aflatoxin biosynthetic genes stands in contrast to the reduced aflatoxin production observed in the Δ*osaA* (Figure 5). Moreover, genes of the cyclopiazonic acid gene clusters were significantly upregulated in Δ*osaA* (Table S3 and Figure S5). The upregulation of CPA genes in Δ*osaA* with respect to the control also contrasts with LC-MS results showing a reduction of CPA in the mutant (Figure 5). Other secondary metabolite gene clusters with significant upregulation included aspirochlorine (8 out of 15 genes, adjusted p-value = 4.8e-5, odds ratio = 22.8) and imizoquins (8 out of 9 genes, adjusted p-value < 1.2e-7, odds ratio = 159.7) (Table S3 and Figure S5).

**Figure 8.**
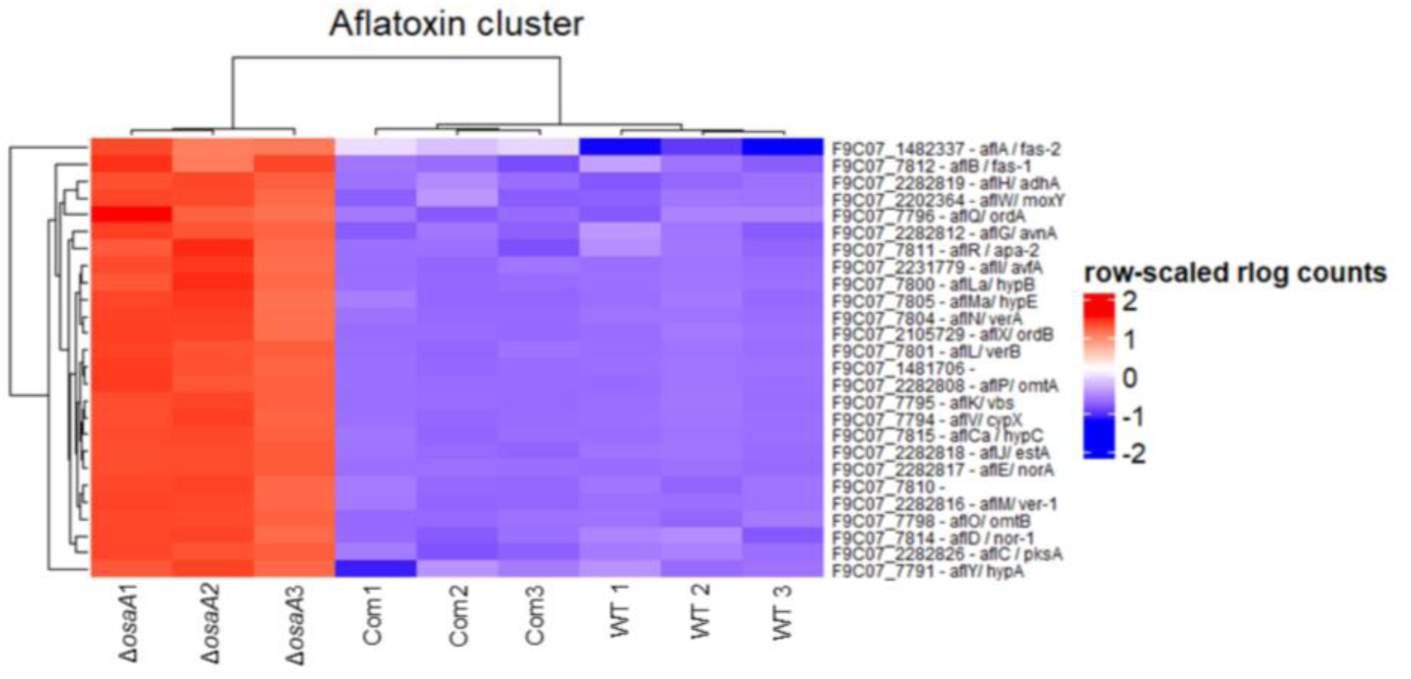
Expression patterns of differentially expressed aflatoxin biosynthesis genes. Hierarchical clustered heatmap displaying row-scaled rlog-transformed read counts for genes in the aflatoxin biosynthesis cluster in wild type (WT), *osaA* deletion and complementation strains. Each row corresponds to an aflatoxin gene, annotated by its locus tag and predicted function. Red indicates higher relative expression, and blue indicates lower relative expression for each gene across samples.

Additional upregulated categories includes monooxygenase activity (GO:0004497; 23 out of 152 genes, adjusted p-value = 3.1e-4, odds ratio = 3.62), phosphopantetheine binding (GO:0031177; 11 out of 41 genes, adjusted p-value = 7.1e-4, odds ratio = 7.3), O-methyltransferase activity (GO:0008171; 9 out of 28 genes, adjusted p-value = 9.3e-4, odds ratio = 9.47), cytochrome P450s (6 out of 20 genes, adjusted p-value = 0.027, odds ratio = 8.5), iron ion binding (GO:0005506; 20 out of 151 genes, adjusted p-value = 0.0059, odds ratio = 3.0), and oxidoreductase activity (GO:0016705; 17 out of 116 genes, adjusted p-value = 0.0059, odds ratio = 3.4), all of which are commonly associated with secondary metabolism (Table S3).

### 2.9. Other osaA-dependent DEGs

#### 2.9.1. Genes involved in transmembrane transporter activity

Transcriptome analysis revealed that *osaA* deletion significantly altered the expression of genes associated with transmembrane transporter activity (Figure S6). In the *osaA* mutant, several trans-membrane transporter genes exhibited marked downregulation while others were upregulated concurrently. These transporters’ sets encompassed diverse families, such as the major facilitator superfamily (MFS), amino acid transporter, and multidrug resistance proteins. Among the downregulated transporters are F9C07_2149740, which encodes an inorganic phosphate transporter; F9C07_2280468, encoding a sugar transporter; F9C07_2282850, predicted to function as a siderochrome–iron transporter; and F9C07_2236497, encoding an MFS multidrug transporter, among others. Those that were upregulated included the MFS transporter (F9C07_2285417), sugar transporter (F9C07_7897), amino acid transporter (F9C07_2232111) and others. These unbalanced transport systems likely have detrimental effects that could affect growth, development, secondary metabolism and virulence.

#### 2.9.2. Genes involved in oxidoreductase activity

RNA-seq analysis also revealed broad transcriptional changes in oxidoreductase-related genes in the absence of *osaA*. Among these, F9C07_2282817, encoding a NADP-dependent oxidoreductase domain-containing protein, was significantly upregulated (Table S2). This enzyme converts NADPH to NADP^+^ which limits the availability of NADPH.

#### 2.9.3. Developmental genes

Transcriptome analysis of the *osaA* mutant revealed differential expressions of key developmental regulators (Table S2). *brlA* (F9C07_2279377) and *flbD* (F9C07_2279158), essential for conidiophore formation, were upregulated in the *osaA* mutant, consistently with the observed increase in conidiation (Figure 3). In contrast, *atfA* (F9C07_2277799), *atfB* (F9C07_9086), *sfgA* (F9C07_10617), *hogA* (F9C07_2156698), *flbC* (F9C07_2282647), and *fluG* (F9C07_5280) were downregulated in the *osaA* mutant. AtfA and AtfB are bZIP transcription factors involved in development and stress response [37], while SfgA is a developmental repressor [38]. HogA also modulates conidiation [39], and FlbC and FluG function upstream as developmental activators [40].

#### 2.9.4. Superoxide dismutase and catalase genes

Transcriptome analysis revealed deletion of *osaA* in *A. flavus* altered the expression of oxidative stress-related genes. Superoxide dismutase was upregulated, whereas catalase was downregulated in the *osaA* mutant. F9C07_2286715, encoding a superoxide dismutase, was upregulated by log₂FC +0.58. On the other hand, F9C07_2285585, F9C07_2284865, F9C07_2237005 and F9C07_2277540, annotated as putative catalase, catalase-like domain-containing protein, heme peroxidase and catalase respectively, were down regulated by log₂FC −1.1, log₂FC −1.4, log₂FC −1.5 and log₂FC −1.6, respectively (Table S2).

#### 2.9.5. chitin synthase gene

Transcriptome analysis also revealed that deletion of *osaA* in *A. flavus* decreased the expression of putative chitin synthase genes. For example, F9C07_366 and F9C07_2286842 annotated as genes encoding chitin synthase and chitin synthesis regulation, respectively, were down regulated by log₂FC −1.2 and log₂FC −2.8 respectively in the *osaA* mutant (Table S2).

### 2.10. Motif enrichment analysis in promoters of differentially expressed genes

To identify potential transcription factor binding sites involved in *osaA*-mediated regulation, we performed motif enrichment analysis on the 1000 bp upstream regions of all DEGs in the Δ*osaA* vs. WT comparison. Using the MEME suite’s Simple Enrichment Analysis (SEA) with the JASPAR 2022 fungi non-redundant database and a p-value threshold of 1e-10, we identified significant enrichment of two similar motifs: MA0382.2 (SKO1 binding motif; consensus sequence DNHDATGACGTAATWDN; p-value = 1.5e-16; enrichment ratio = 1.5) and MA0286.1 (CST6 binding motif; consensus sequence RTGACGTMA; p-value = 6.6e-14; enrichment ratio = 2.3) (Figure 9). Among the DEGs, 347 contained the SKO1 motif and 110 contained the CST6 motif, with 112 genes containing both motifs.

**Figure 9.**
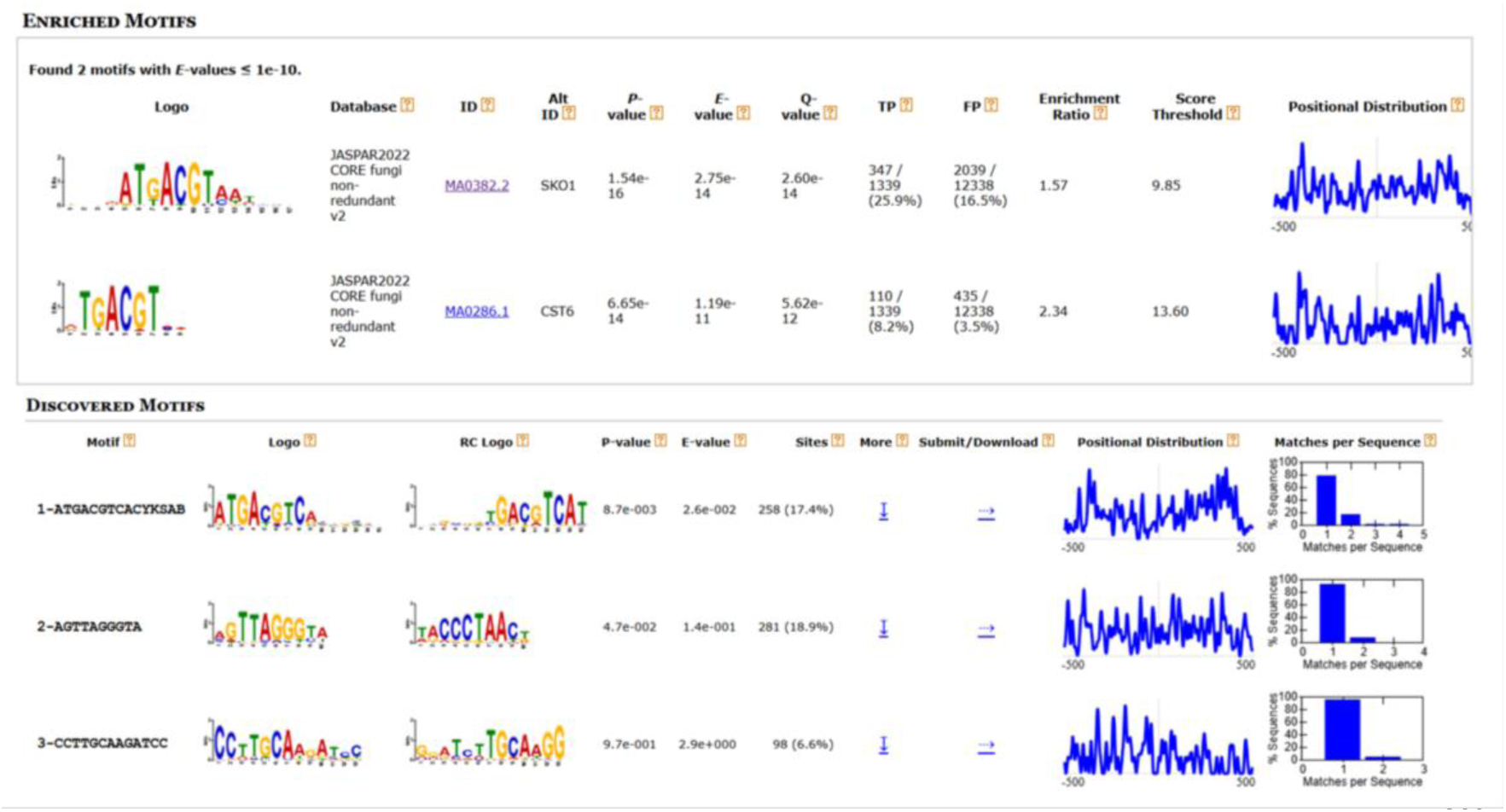
Motif enrichment analysis of promoters of *osaA*-dependent genes. Motif enrichment using Simple Enrichment Analysis (SEA) from the MEME suite with the JASPAR 2022 fungi non-redundant database identified two similar significantly enriched motifs: the SKO1 binding motif (MA0382.2) and the CST6 binding motif (MA0286.1). De Novo motif discovery using STREME identified three additional significant motifs, including one resembling the SKO1 binding site (MA0382.2), one with similarity to MA0403.2, and a novel motif of unknown function.

Additionally, de novo motif discovery using STREME identified three significant motifs in the promoters of DEGs. The first motif closely resembled the SKO1 binding site (MA0382.2) identified with SEA, while the second showed similarity to the MA0403.2 motif in the JASPAR database (Figure 9). The third motif did not match any known fungal transcription factor binding sites, potentially representing a novel regulatory element associated with *osaA*-mediated gene expression.

## 3. Discussion

Novel regulatory genes, and their products, that govern fungal development, secondary metabolism and pathogenesis could serve as targets for control strategies to effectively decrease *A. flavus* colonization and aflatoxin contamination. Gti1/Pac2 (WOPR) domain proteins represent a class of fungal-specific transcription factors involved in regulating those cellular processes [24]. In our study, we identified that in the agriculturally important fungus *A. flavus,* OsaA contains a typical WOPR domain in its N-terminal region, composed of two conserved subdomains separated by a short linker. Our results also support that the *Aspergillus* OsaA protein is phylogenetically related to the Gti1/Wor1/Ryp1/Mit1/Rge1/Ros1/Sge1 proteins described in other fungal species [26]. Structural analysis of another study shows WOPR subdomains are tightly connected by a β-sheet, with the linker extending outward from the DNA, and the β-strands of each subdomain interdigitating to form a stable structure [24].

In studies using the model filamentous fungus *A. nidulans*, the *osaA* homolog has been shown to regulate development by repressing sexual reproduction downstream of the global regulator VeA [33]. Similarly, *osaA* plays a key role in *A. fumigatus* morphology. Absence of *osaA* significantly reduces colony growth, germination rate, conidial production in this opportunistic human pathogen [34]. In contrast, our study revealed that loss of *osaA* in *A. flavus* leads to increased conidial production. Consistent with these phenotypes, transcriptome data revealed upregulation of key conidiophore developmental genes in the *osaA* mutant. Notably, *brlA* (F9C07_2279377) and *flbD* (F9C07_ 2279158) which are central developmental regulators, were upregulated. BrlA, is a C_2_H_2_ zinc finger transcription factor, essential for initiating conidiophore formation, while FlbD acts upstream to activate *brlA* expression [37,41]. These gene expression patterns support the observed increase in conidiation in the *osaA* mutant.

In addition to conidiation, *osaA* influences sclerotial formation in *A. flavus. osaA* is indispensable for normal sclerotial production rates. Colony of the *osaA* deletion strain produced significantly fewer sclerotia. This effect was more dramatic on top-agar inoculated cultures where sclerotial formation is completely abolished in the absence of *osaA*. This dramatic reduction may be due to high spore density, and therefore a crowdedness effect in the latter cultures, leading to an accentuated competition. In addition, our transcriptome data showed that transcription factor genes *atfA (F9C07_2277799)* and *atfB (F9C07_9086)* were downregulated in the *osaA* mutant. These genes positively affect sclerotial formation [42], therefore, a reduction of *atfA* and *atfB* expression in the *osaA* deletion mutant could contribute the observed decrease in sclerotial production.

Since development and secondary metabolism are found to be genetically linked in *Aspergillus* species [5], we also investigated the possible role of *osaA* in regulating production of natural compounds in *A. flavus*, particularly mycotoxins. Our findings demonstrate that *osaA* positively regulates the biosynthesis of both AFB_1,_ and CPA in *A. flavus*. Unexpectedly, most aflatoxin biosynthetic genes, and genes of CPA gene clusters, were upregulated in the *osaA* mutant compared to the control. This contrasts with chemical analyses results, which showed a clear reduction in AF, as well as CPA production, in the mutant. This discrepancy suggests the presence of a feedback mechanism, where the fungal system attempts to compensate for reduced mycotoxin levels by up-regulating gene expression. The deletion of *osaA* may impair post-transcriptional regulation, preventing proper translation, processing, or localization of AF biosynthetic enzymes. Aflatoxin biosynthesis and transport are known to involve specialized vesicles called aflatoxisomes [43]. In *Aspergillus parasiticus*, the vesicle-vacuole system is essential for converting sterigmatocystin (ST) to aflatoxin B_1_ and compartmentalizing the toxin [44]. Key enzymes such as Nor-1, Ver-1, and OmtA are synthesized in the cytoplasm and transported to vacuoles via the cytoplasm-to-vacuole targeting (Cvt) pathway [44]. More recently, lipid droplets have also been implicated in aflatoxin biosynthesis and export [45]. Given the connection between aflatoxisome function and membrane transport, defects in transporter gene expression in the *osaA* mutant may further impair AF production. Expression of several transmembrane transporter genes were altered in the *osaA* mutant. This widespread changes in transporters could restrict the availability of key nutrients and cofactors necessary for aflatoxin biosynthesis, including phosphate, iron ions, and sugar. Furthermore, aflatoxin biosynthesis requires NADPH as a cofactor, especially for the cytochrome P450 monooxygenase OrdA, which catalyzes the conversion of O-methylsterigmatocystin (OMST) and dihydro-OMST into aflatoxins B_1_, B_2_, G_1_, and G_2_ [46]. In the *osaA* mutant, we observed upregulation of a gene *(F9C07_2282817*) encoding NADP-dependent oxidoreductase domain-containing protein (Table S2), which likely catalyze the conversion of NADPH to NADP⁺. Thereby decreasing the availability of NADPH required for aflatoxin biosynthetic enzymes which would further contributing to the observed reduction in AF production.

Additionally, downregulation of *atfA* (F9C07_2277799) and *atfB* (F9C07_9086) may partially explain the outcome of reduction of aflatoxin production despite the upregulation of aflatoxin biosynthetic genes. The bZIP transcription factors were demonstrated as co-regulators of aflatoxin biosynthesis and oxidative stress in *A. parasiticus* [47,48]. The reduced expression of bZIP transcription factors may compromise the fungus’ ability to manage oxidative stress, which in turn can negatively influence aflatoxin biosynthesis.

Expression of other secondary metabolite genes, including genes in the imizoquin and aspirochlorine biosynthetic clusters, were upregulated by *osaA*. Imizoquin possesses reactive oxygen species (ROS)-quenching properties which help to maintain ROS homeostasis and promote spore germination in the *A. flavus* [49]. In *A. flavus*, the imizoquin biosynthetic pathway is enhanced by specific oxylipins, contributing to the inhibition of other metabolic pathways such as aflatoxin biosynthesis [50]. Additionally, genes within the aspirochlorine biosynthetic cluster were also upregulated in the *osaA* mutant. Aspirochlorine is a halogenated epidithiodiketopiperazine mycotoxin with antibiotic activity, also known as antibiotic A30641[51]. The increased expression of this cluster in the *osaA* mutant raises the possibility that *osaA* could contribute to antifungal defense and may represent a promising target for antifungal drug development.

Production of secondary metabolites, including mycotoxins, is also influenced by environmental stressors. *A. flavus* withstands various biotic and abiotic stresses, including fluctuations in temperature and oxidative stress. This fungus can grow and produce aflatoxins over a wide temperature range, with thermal tolerance being an important factor influencing its pathogenicity and toxin biosynthesis [1,52]. As mentioned above, oxidative stress also plays a critical role in regulating aflatoxin production, as *A. flavus* responds to reactive oxygen species through complex regulatory pathways that modulate secondary metabolite biosynthesis [1,53]. The possible role of *osaA* in *A. flavus* in oxidative stress response were evaluated in our study. The *osaA* mutant showed increased sensitivity to hydrogen peroxide. However, this strain presented greater resistance to menadione compared to control strains. In *Saccharomyces cerevisiae*, *rad9* mutants showed normal resistance to menadione but were up to 100-fold more sensitive to H₂O₂ [54], indicating distinct features in the cellular responses [55]. Menadione generates superoxide radicals, which can cause oxidative damage. If the mutation enhances superoxide dismutase activity, the mutant could exhibit increased resistance to menadione. However, in the case of hydrogen peroxide, its detoxification relies on catalases [56–58]. Our transcriptome indicated that the catalase gene (F9C07_2237005), was significantly down regulated in the *osaA* mutant, which could contribute to the increase sensitivity to hydrogen peroxide in this strain. Conversely, superoxide dismutase gene (F9C07_2286715) expression was slightly upregulated, and could contribute to the increased resistance to menadione observed in the *osaA* mutant. Several studies have demonstrated that certain aflatoxin inhibitors reduce AFB_1_ production by modulating antioxidant activity. For example, ascorbic acid and cinnamaldehyde significantly decreased AFB_1_ levels while increasing superoxide dismutase activity [47,53,59]. Enhanced superoxide dismutase activity in Δ*osaA* could contribute to a reduction in aflatoxin biosynthesis. In addition, downregulation of Pentose Phosphate Pathway (PPP) genes in *osaA* mutant can also weaken both oxidative stress tolerance. The PPP is crucial for producing NADPH, which acts as a reducing equivalent to maintain the redox balance inside fungal cells. NADPH is required for the function of antioxidant enzymes such as catalase. When PPP genes are downregulated, the NADPH pool decreases, leading to increased sensitivity to reactive oxygen species (ROS) [60]. Since *osaA* mutant presents downregulation of PPP genes, it is likely that this strain is unable to efficiently neutralize ROS.

In addition to oxidate stress, we also assessed *osaA*-dependent temperature sensitivity. Both low or high temperatures can affect *A. flavus* growth [52,61]. We found that the decrease in growth rate in the *A. flavus osaA* mutant compared to the wild type was more pronounced at lower temperature (25°C) compared to that observed at 30°C. All strains exhibited growth at 42°C. This contrasts with findings in *A. fumigatus*, where *osaA* is specifically required for growth at elevated temperatures (42°C) [34].

The changes in sensitivity to environmental stresses could also be influenced by alterations in the fungal cell wall [62]. The fungal cell wall is a rigid, dynamic structure essential for fungal viability [63]. In *A. fumigatus*, deletion of *osaA* compromises cell wall stability, further implicating its role in stress adaptation [34]. Thus, the increase in cold sensitivity observed in the *A. flavus* Δ*osaA* strain could be the result of possible changes in its cell wall composition and integrity. Our study revealed a significant reduction in chitin content in the *osaA* mutant with respect to the wild type, which suggests that *osaA* positively regulates chitin synthesis in *A. flavus*. Furthermore, the chitin synthase gene (F9C07_366) is downregulated in the *osaA* mutant, which could contribute to the decreased chitin level in this strain’s cell wall.

The decrease in chitin in the absence of *osaA*, and possible concomitant reduction in cell wall integrity, together with the additional alterations in biological processes mentioned above, could contribute to a reduction in virulence. We investigated the role of *osaA* in seed infection. Our results show that *osaA* is require for wild-type levels of seed colonization, as its deletion reduces fungal burden in corn kernels. Our transcriptome analysis indicated down regulation of apoplast, apoplastic effectors and cytoplasmic effector genes in the Δ*osaA*, which could lead to reduce pathogenicity in seed [64,65]. Also, the downregulation of pentose phosphate pathway genes in the *osaA* mutant and its effect on oxidative stress could lead to impaired infection ability and attenuated virulence. Importantly, the absence of *osaA* led to a marked decrease in AFB_1_ and AFB_2_ production also during infected seeds.

To further gain insight into the *osaA*’s regulatory role, a promoter analysis was conducted using known fungal transcription factors binding motifs from the JASPAR database. Two motifs were MA0382.2 (SKO1 binding motif) and MA0286.1 (CST6 binding motif) which were identified by motif enrichment analysis on the 1000 bp upstream regions of all DEGs in the Δ*osaA* vs. wild type comparison. The SKO1 protein that is associated with the MA0382.2 motif seems to operate in response to osmotic stress and cell wall damage. In *Aspergillus cristatus* the SKO1 protein, a homolog of the yeast SKO1p, functions as a transcription factor involved in regulating osmotic stress response, similar to its role in *S. cerevisiae* [66]. This further suggests that OsaA, perhaps in cooperative with a SKO1 homolog in *A. flavus*, may regulate stress-responsive pathways via transcriptional control. With respect to CST6, a recent study has demonstrated that this transcription factor is involved in azole susceptibility in *Candida glabrata,* showing that CST6 regulates a broad gene-network, including genes involved in respiration, cell-wall and adhesins in *C. glabrata.* It is possible that OsaA could also be associated with antifungal drug susceptibility in *A. flavus* [67]. Future studies will focus on possible interactions between these relevant regulatory factors.

In summary, *osaA* functions as a central regulator integrating fungal development, environmental stress management, secondary metabolite production, and virulence in *A. flavus*. These findings provide valuable insights into the regulatory networks underlying fungal pathogenicity and identify *osaA* as a promising target for future antifungal strategies.

## 4. Materials and Methods

### 4.1. Sequence Analysis

The deduced amino acid and nucleotide sequences used in the phylogenic analysis were obtained from FungiDB (https://fungidb.org/fungidb/app/; version: Release 68, 7 May 2024). All gene and protein accessions are listed in Table 1.

**Table 1.**
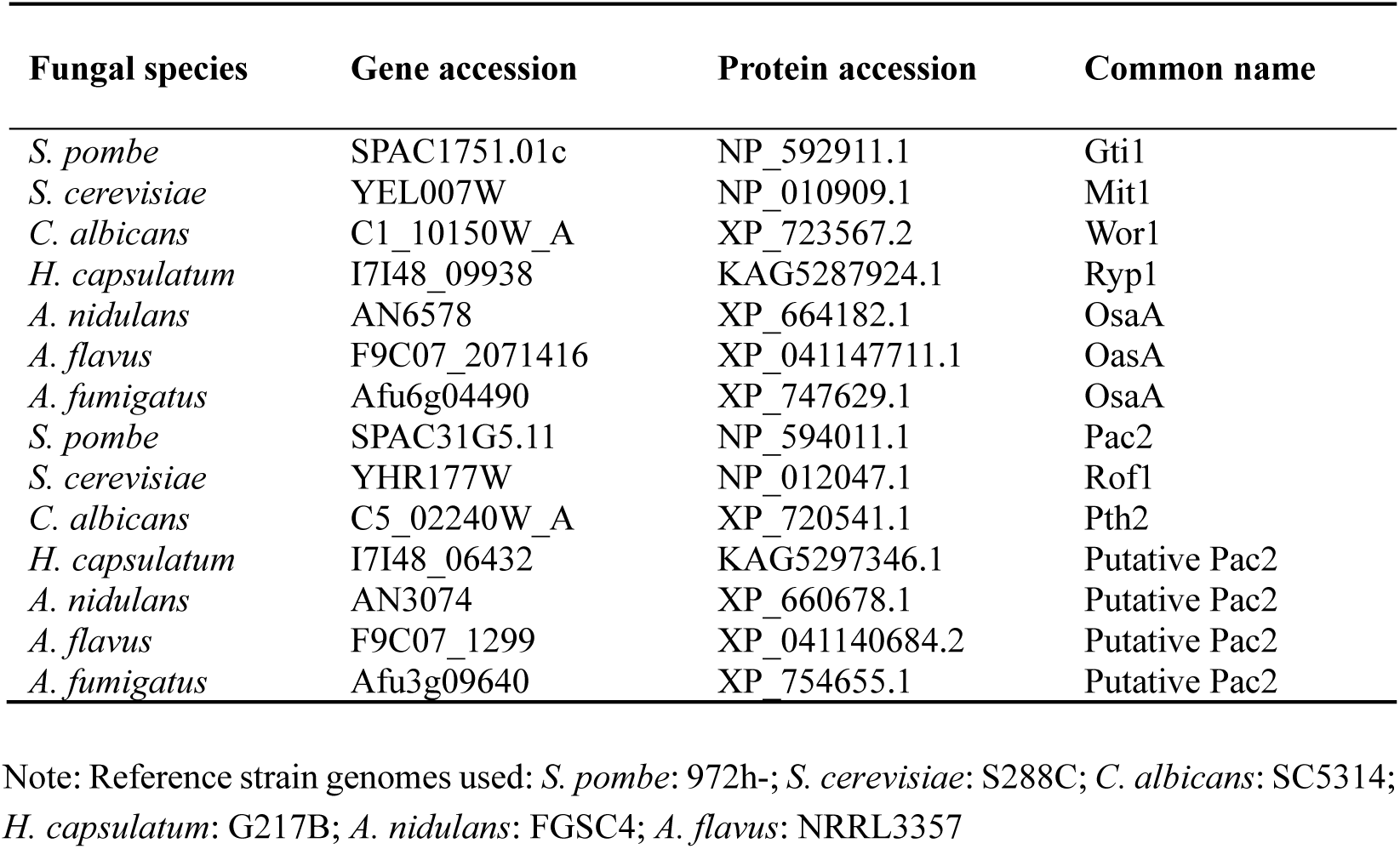
Accession for genes/proteins used in phylogenetic analysis.

Corresponding protein accession from NCBI were found by conducting a blastP analysis using sequences obtained from FungiDB. Using Geneious software, a MAFFT multisequence alignment (Version1.2.2) was conducted using the default parameters to align the protein sequences [68]. Subsequently, a maximum-likelihood tree was conducted using PhyML (version 3.3.20180621) with a LG substitution model and 1,000 bootstrap replicates [69].

### 4.2. Strains and culture conditions

The *A. flavus* strains utilized in this study were the CA14 *pyrG*-1control strain, *osaA* deletion strain (TMR1.1) and *osaA* complementation strain (TFEH 8.1) (Table 2). All strains were grown on commercial PDA (Potato Dextrose Agar) medium (BD Difco™) unless otherwise indicated. Fungal strains were maintained in 30% glycerol stocks at −80°C.

**Table 2.** Strains used in this study.

| Strain name | Genotype | Source |
| --- | --- | --- |
| CA14_pSL82 | $\Delta pyrG, niaD^+, ptrA^S, \Delta ku70$ | Gift from Dr. Jeffrey Cary |
| CA14 *pyrG*-1 (WT) | $pyrG^+, niaD^+, ptrA^R, \Delta ku70$ | Gift from Dr. Jeffrey Cary |
| TMR1.1 | $\Delta pyrG, \Delta osaA::pyrG^{A. fumigatus}, niaD^+, ptrA^S, \Delta ku70$ | This study |
| TFEH8.1 | $\Delta osaA::pyrG, osaA::trpC::ptrA^R::pyrG^{A. fumigatus}, niaD^+, \Delta ku70$ | This study |

### 4.3. Generation of the osaA deletion strain

The Δ*osaA* strain was generated by a gene replacement strategy described by [35]. The deletion DNA cassette was generated by fusion PCR. The 5′ untranslated region (1.854 kb) and 3′ untranslated region (1.778 kb) flanking the *osaA* coding sequence were amplified from *A. flavus* genomic DNA using primer pairs 3011/3012 and 3013/3014, respectively (Table S1). The *A. fumigatus pyrG* selection marker was amplified from plasmid p1439 [70] with primers 3015 and 3016. These three fragments were fused via PCR using primers 3017 and 3018 to generate the knockout cassette. The construct was introduced into *A. flavus* CA14_pSL82 (*pyrG-, niaD+*, *ptrA^S^*, Δ*ku70*) protoplasts via polyethylene glycol-mediated transformation. Successful deletion of the *osaA* coding region and replacement with *pyrG* was confirmed by diagnostic PCR using primers 3011 and 2846, resulting in the Δ*osaA* strain (TMR1.1).

### 4.4. Generation of the osaA complementation strain

The complementation strain (TFEH8.1) was constructed by re-introducing the wild type *osaA* allele into the Δ*osaA* strain at the same locus. To generate the complementation cassette, four DNA fragments were PCR-amplified: the *osaA* coding sequence along with a 2 kb upstream region (5′UTR) using primers 3065 and 3066; the *trpC* terminator using primers 2869 and 2870; the *ptrA* selection marker using primers 3067 and 3068; and the *A. fumigatus pyrG* gene from plasmid p1439 using primers 2871 and 2872. These four fragments were fused in a single PCR reaction using primers 3063 and 3064. The resulting fusion cassette was introduced into the Δ*osaA* strain TMR1.1 via polyethylene glycol-mediated transformation [35]. Transformants were screened and verified by diagnostic PCR using primers 3063 and 3047. The selected transformant was designated as TFEH8.1.

### 4.5. Morphological analysis

#### 4.5.1. Colony growth

To assess the role of *osaA* in vegetative colony growth, *A. flavus* CA14 wild type, Δ*osaA* and *osaA* complementation strain were point-inoculated onto commercial PDA medium (BD Difco™) and incubated at 30°C. Colony growth was measured as colony diameter at both 5-and 7-days post-inoculation. All experiments were performed in triplicate.

#### 4.5.2. Conidial production

To investigate the role of *osaA* in conidiation, *A. flavus* CA14 wild type, Δ*osaA* and *osaA* complementation strain were point-inoculated onto PDA medium and incubated at 30°C in the dark. Conidiation was evaluated at 5- and 7-days post-inoculation. Agar cores (10 mm in diameter) were collected from regions 5 mm and 20 mm away from the colony center. The samples were homogenized in sterile water, and conidia were quantified using a haemocytometer (Hausser Scientific, Horsham, PA, USA) under bright-field microscopy. Additionally, close-up images of conidiophores from both regions were captured using a Leica dissecting microscope at 32x magnification.

#### 4.5.3. Sclerotial development

To determine the role of *osaA* in sclerotial production, *A. flavus* CA14 wild type, Δ*osaA* and *osaA* complementation strain were both point-inoculated and top-agar inoculated (10⁶ spores/mL) onto Wickerham agar medium (Per liter: 2.0 g yeast extract, 3.0 g peptone, 5.0 g corn steep solids, 2.0 g dextrose, 30.0 g sucrose, 2.0 g NaNO_3_, 1.0 g K_2_HPO_4_·3H_2_O, 0.5 g MgSO_4_·7H2O, 0.2 g KCl, 0.1 g FeSO_4_·7H_2_O (10-fold the original recipe), and 15.0 g agar per liter [pH 5.5]). Cultures were incubated at 30°C in the dark for 14 days. Following incubation, cultures were washed with 70% ethanol to remove conidia and enhance sclerotial visibility. Micrographs were captured using a Leica MZ75 dissecting microscope equipped with a DC50LP camera (Leica Microsystems, Inc., Buffalo Grove, IL, USA) at 12.5x and 32x magnifications.

### 4.6. Aflatoxin B_1_ production analysis

To evaluate whether *osaA* regulates AFB_1_ biosynthesis, *A. flavus* CA14 wild type, Δ*osaA* and *osaA* complementation strain were point-inoculated on PDA medium and incubated in the dark at 30°C for 7 days. Three 16-mm-diameter cores were collected approximately 5 mm from the colony center. AFB_1_ was extracted using 5 mL of chloroform per sample. The extracts were evaporated to dryness and subsequently resuspended in 200 µL of chloroform. AFB_1_ detection was performed via thin-layer chromatography (TLC) using silica-precoated Polygram Sil G/UV254 TLC plates (Macherey-Nagel, Bethlehem, PA, USA) and a solvent system composed of toluene:ethyl acetate:formic acid (50:40:10). After air-drying, the TLC plates were sprayed with a 12.5% aluminum chloride (AlCl₃) solution in ethanol, baked at 80°C for 10 minutes, and visualized under UV light at 375 nm. Aflatoxin B_1_ standard was obtained from Sigma-Aldrich (St. Louis, MO, USA). Samples were also analyzed to identify other secondary metabolites by LC-MS (section 4.7) and to further quantify Aflatoxin B_1_ by UPLC with fluorescence detection (section 4.10.2).

### 4.7. Secondary metabolite analysis by LC-MS

*A. flavus* secondary metabolites in the extracts were analyzed on a Waters Acquity UPLC system coupled to a Waters Xevo G2 XS QTOF mass spectrometer. Extract injections (1 µl) were separated on a Waters BEH C18 1.7 µm, 2.1 x 50 mm column with the following gradient solvent system: (0.5 ml/min, solvent A: 0.1% formic acid in water; solvent B: 0.1% formic acid in acetonitrile): 5% B (0-1.25 min.), gradient to 25% B (1.25-1.5 min.), gradient to 100% B (1.5-5.0 min.), 100% B (5.0-7.5 min.), then column equilibration to 5% B (7.6-10.1 min.). The Z-spray ionization source was run in ESI+ mode using MassLynx 4.2 software with the following settings: source temperature: 100 °C, desolvation temperature: 250 °C, desolvation gas flow: 600 L/h, cone gas flow: 50 L/h, capillary voltage: 3.0 kV, sampling cone voltage: 40 V. Analyses were performed in sensitivity and continuum mode, with a mass range of m/z 50–1200 and a scan time of 0.1 s. A data-independent acquisition method with elevated collision energy (MS^E^) was used with 6 eV low energy and a high energy ramp from 15−45 eV. Mass data were collected from 2.0-6.0 min. then imported, analyzed, and quantified on Waters UNIFI 1.9.4 software using “Quantify Assay Tof 2D” analysis method with lock mass corrected by UNIFI. Aflatoxin B_1_ and CPA standards were purchased from Sigma-Aldrich (St. Louis, MO, United States). Metabolite content is expressed in ppb (ng/g samples).

### 4.8. Environmental stress tests

#### 4.8.1. Temperature sensitivity

To assess the role of *osaA* in temperature sensitivity, *A. flavus* CA14 wild type, Δ*osaA* and *osaA* complementation strain were point-inoculated on PDA medium and incubated in the dark at 25°C, 30°C, 37°C, and 42°C for 5 days. Colony diameters were measured, and growth was assessed across three biological replicates. The percentage reduction in growth was calculated by comparing colony diameters at 30°C to those at 25°C, 37°C, and 42°C.

#### 4.8.2. Oxidative stress sensitivity

To evaluate the potential role of *osaA* in oxidative stress sensitivity, *A. flavus* CA14 wild type, Δ*osaA* and complementation strain were point-inoculated on PDA plates supplemented with 0.4, 0.5, and 0.6 mM menadione and incubated at 30°C in the dark for 3 days. Colony growth was measured, and the percentage of growth reduction was calculated relative to control plates without menadione. Experiments were performed in triplicate. Additionally, oxidative stress sensitivity was tested using hydrogen peroxide. The same strains were point-inoculated on PDA medium supplemented with 0.1%, 0.2%, 0.3%, and 0.4% hydrogen peroxide and incubated at 30°C for 3 days. Four biological replicates were included. Growth inhibition was calculated by comparing colony diameters from treated versus untreated conditions.

### 4.9. Chitin analysis

To measure the levels of chitin in the *A. flavus* CA14 wild type, Δ*osaA* and *osaA* complementation strain cell wall, a previously described protocol [71] was followed with minor modifications. Briefly, *A. flavus* strains were inoculated into 50 mL of liquid commercial PDB (BD Difco™) medium (10⁶ spores/mL) and incubated at 30°C for 48 h at 250 rpm. Mycelia were harvested using Miracloth (Calbiochem, San Diego, CA), washed three times with sterile distilled water, and stored at −20°C.

For cell wall analysis, frozen mycelia were resuspended in 1 mL of cell wall buffer (2% SDS in 50 mM Tris-HCl, pH 7.5, supplemented with 100 mM Na-EDTA, 40 mM β-mercaptoethanol, and 1 mM PMSF) and boiled for 15 min to remove unbound proteins and soluble sugars. After boiling, samples were washed three times with sterile ddH₂O and lyophilized overnight.

Approximately 40 mg of lyophilized mycelia per strain (three replicates each) were treated with 3% NaOH at 75°C for 1 h. Samples were centrifuged at 15,000 g for 15 min. The remaining pellet was digested with 96% formic acid at 100°C for 4 h. After evaporation of formic acid, residues were resuspended in 1 mL of sterile ddH₂O for chitin.

Quantification of chitin was performed as described in [72] and measured absorbance at 520 nm using an Epoch spectrophotometer (BioTek, Winooski, VT).

### 4.10. Virulence studies by seed infection assay

#### 4.10.1. Kernel screening assay

A Kernel Screening Assay (KSA) was performed on corn seeds from the aflatoxin sensitive B73 corn line as a plant model as previously described in [73]. Conidiospores from the CA14 wild-type control, Δ*osaA* and *osaA* complementation strain cultivated on 2x V8 agar (100 mL V8 juice, 40 g agar, per liter of medium and pH 5.2) and incubated at 30°C in the light for 7 days. Spores were collected in 25 mL of water in aseptic conditions and quantified using an Olympus Automated Cell Counter Model R1 (Olympus Corporation, Shinjuku, Tokyo, Japan). Undamaged B73 seeds of a relatively similar size were collected and surface sterilized using 10% bleach. The seeds were infected with spores of the *A. flavus* strains by being soaked in 10 mL of 1.0 x 10^4^ spores/mL spore suspension for each strain. Uninfected B73 seeds, serving as control (mock), were also treated with water only, in a manner similar to the seeds infected with *A. flavus* spores. The falcon tubes containing the seeds and spore suspensions were rocked for 3 minutes to ensure equal distribution of inoculum throughout the seeds. The inoculum was drained from the falcon tubes and the seeds transferred to sterile petri plate lids (60 x 15 mm). These were placed onto larger trays containing 3 MM Whatman filter paper and 50 mL of sterile water reservoir to provide humidity. The experiment was done in replicates of six for each strain. The cultures were then incubated at 30°C under dark conditions for 7 days. The seeds were then photographed and harvested by flash freezing the seeds in liquid nitrogen. Prior to LC-MS analysis the seeds were pulverized using a SPEX SamplePrep 2010 Geno/Grinder (SPEX SamplePrep, Metuchen, NJ, USA) prior to lyophilization and storage at −80°C.

#### 4.10.2. Extraction and aflatoxin analysis from seeds

Lyophilized ground corn powder (100 mg) was transferred into a 2 mL Eppendorf tube with methanol (1 mL) and vortexed vigorously for 15 sec. Tubes were secured onto an orbital platform shaker (Solaris 2000, Thermo Fisher Scientific) then rotated at 200 rpm, at RT and in darkness, approximately 22 h. Next, tubes were centrifuged for 5 min at 14,000 rpm to remove particulate, and a portion of the particulate-free extract (1 mL) was transferred to a new tube for storage and analysis. The aflatoxin-containing solution was analyzed (1 µl injections) using a Waters ACQUITY UPLC system (40% methanol in water, BEH C18 1.7 μm, 2.1 mm × 50 mm column) with fluorescence detection (Ex = 365 nm, Em = 440 nm). Samples were diluted if the aflatoxin signal saturated the detector. Analytical standards (Sigma-Aldrich, St. Louis, MO, United States) were used to identify and quantify aflatoxins AFB_1_ and AFB_2_. Aflatoxin content was expressed in ppb (ng/g corn dry weight).

#### 4.10.3. Ergosterol extraction and analysis

Lyophilized ground corn (50 mg) was transferred to 15 ml Falcon tubes. Alcoholic potassium hydroxide (KOH) was prepared: 25 g KOH was dissolved in 35 mL water then 100% ethanol was added for total volume of 100 mL. 3 ml of the alcoholic KOH solution was added to each sample tube, the mixture was vortexed for 1 min, and then incubated at 85 °C in a water bath for 1.5 h. The samples were then cooled to room temperature, distilled water (1 mL) and hexanes (3 mL) were added to the tubes and vortexed vigorously for 3 min to extract ergosterol. The hexanes layers were carefully transferred to clean 4 ml glass vials and concentrated via SpeedVac (Savant, Thermo Scientific). The concentrated extracts were redissolved in methanol (500 µl) for analysis. The redissolved solution was analyzed (1 µl injections) using a Waters ACQUITY UPLC system (95% methanol in water, BEH C18 1.7 μm, 2.1 mm × 50 mm column) with UV detection (λ = 282 nm). An analytical standard of ergosterol (Sigma-Aldrich, St. Louis, MO, United States) was used for quantification. Ergosterol content is reported as µg/g corn dry weight.

### 4.11. osaA expression analysis

Plates containing 25 mL of PDB were inoculated with 10^6^ spores/mL of *A. flavus* CA14 wild type, Δ*osaA* and *osaA* complementation strain, and incubated in the dark at 30°C, for 72 h. Total RNA was extracted from lyophilized mycelial samples using TRIsure™ (Meridian Bioscience, Bioline, Cincinnati, OH, USA) according to the manufacturer’s instructions. For gene expression analysis, 1µg of total RNA was treated with the Ambion TURBO DNA-free™ Kit (Thermo Fisher Scientific, Waltham, MA, USA) to remove genomic DNA. First-strand cDNA was synthesized using Moloney murine leukemia virus (MMLV) reverse transcriptase (Promega, Madison, WI, USA).

Quantitative real-time PCR (qRT-PCR) was carried out using either the Bio-Rad CFX96 Real-Time PCR System or the Applied Biosystems 7000 system. Reactions were prepared using either iQ SYBR Green Supermix (Bio-Rad, Hercules, CA, USA) or SYBR Green JumpStart Taq ReadyMix (Sigma-Aldrich, St. Louis, MO, USA) for fluorescence detection. The primer pairs 3071 and 3072 were used for target gene amplification. Expression levels were normalized to 18S rRNA as an internal control. Relative expression was calculated using the 2^−ΔΔ*CT*^ method [74].

### 4.12. Transcriptome analysis

#### 4.12.1. RNA Extraction

Plates containing 25 mL of liquid potato dextrose broth (PDB) were inoculated with 10⁶ spores/mL of *A. flavus* CA14 wild type, Δ*osaA* and *osaA* complementation strain incubated in the dark at 30°C for 3 days. Following incubation, mycelia was harvested, frozen in liquid nitrogen, and lyophilized. Total RNA was extracted from the lyophilized mycelia using the RNeasy Plant Mini Kit (Qiagen, Germantown, MD, USA) according to the manufacturer’s instructions.

#### 4.12.2. RNA sequencing

RNA sequencing generated an average of 56 million 150 bp paired-end reads per sample. Adapters and low-quality sequence were removed from the reads using fastp [75]. Trimmed reads were aligned to the *A. flavus* NRRL 3357 genome (GCA_009017415.1) using STAR (version 2.7.10b) with the following settings “--alignIntronMax 1000--twopassMode Basic--quantMode Gene-Counts”. The forward-stranded gene-level read counts output by STAR were used as input for differential expression analysis. Differentially expressed genes (DEGs) between the samples were identified using DESeq2 (version 1.36.0) [76]. Log2 fold changes and p-values were estimated using the lfcShrink function with type=“ashr” [77] and alpha=0.05 Results were filtered to retain genes with adjusted p-value < 0.05 and absolute log2 fold change > 1. Functional enrichment analysis of DEGs was performed using the enrichment function in the BC3NET R package [78] which implements a one-sided Fisher’s exact test. False discovery rate was controlled using the p.adjust R function with method=“fdr” [79]. Heatmaps were created using the tidyHeatmap R package [80] with regularized log-transformed counts and row scaling enabled. All heatmaps display only genes differentially expressed in the deletion vs. wild-type comparison.

Transcription factor binding site enrichment in promoter sequences was analyzed using the MEME Suite (version 5.5.7) [81]. The sequence of 1,000 base pairs upstream of the coding start site of each DEG was used as the input and the corresponding upstream region from all other genes was used as the background. Known transcription factors from the JASPAR non-redundant fungi 2022 database were used with the Simple Enrichment Tool from MEME with a e-value threshold of 1e-10. De Novo motif analysis was also performed using STREME from the MEME suite with default options, the same input and background sequences as above, and an e-value threshold of 1e-10. Motifs discovered using STREME were compared to known motifs using the Tomtom tool in MEME.

### 4.13 Statistical analysis

Statistical analysis was applied to analyze all quantitative data in this study utilizing analysis of variance (ANOVA) in conjunction with a Tukey multiple-comparison test using a *p* value of *<*0.05 for samples that are determined to be significantly different unless otherwise indicated.

## Supplementary Materials

Figure S1: Sequence alignment of Gti1/Pac2 orthologs; Figure S2: Role of *osaA* in temperature sensitivity in *A. flavus*; Figure S3: Role of *osaA* in oxidative stress sensitivity in *A. flavus*; Figure S4: Role of *osaA* in the synthesis of chitin in *A. flavus*; Figure S5: Differential expression of genes from the cyclopiazonic acid gene clusters, imizoquin genes and aspirochlorine gene; Figure S6. Differential expression of transmembrane transporter genes; Table S1: Primers used in this study; Table S2: All differentially expressed genes (DEG) and their log2 fold change. Table S3: Enriched annotation terms in differentially expressed genes in the *osaA* deletion vs wild type comparison.

## Author Contributions

Conceptualization: F.E.H. and A.M.C.; Methodology: F.E.H., A.D., J.M.L. M.D.L., B.M.M. and A.M.C.; Software: B.M.M.; Validation: F.E.H., A.D., J.M.L. M.D.L., B.M.M. and A.M.C.; Formal Analysis: F.E.H., A.D., J.M.L. M.D.L., B.M.M. and A.M.C.; Investigation: F.E.H., A.D., J.M.L. M.D.L., B.M.M. and A.M.C.; Resources: A.M.C.; Data Curation: F.E.H., A.D., J.M.L. M.D.L., B.M.M. and A.M.C.; Writing – Original Draft Preparation: F.E.H. and A.M.C.; Writing – Review & Editing: F.E.H. and A.M.C.; Visualization: F.E.H. and A.M.C.; Supervision: A.M.C.; Project Administration: A.M.C.; Funding Acquisition: A.M.C.

## Institutional Review Board Statement

Not applicable

## Informed Consent Statement

Not applicable

## Data Availability Statement

None.

## Acknowledgments

We would like to thank Scott Grayburn, Makayla R. Ritko and Christopher Salgado for their technical support.

## Disclaimer/Publisher’s Note

The statements, opinions and data contained in all publications are solely those of the individual author(s) and contributor(s) and not of MDPI and/or the editor(s). MDPI and/or the editor(s) disclaim responsibility for any injury to people or property resulting from any ideas, methods, instructions or products referred to in the content.

